# P-sort: an open-source software for cerebellar neurophysiology

**DOI:** 10.1101/2021.03.16.435644

**Authors:** Ehsan Sedaghat-Nejad, Mohammad Amin Fakharian, Jay Pi, Paul Hage, Yoshiko Kojima, Robi Soetedjo, Shogo Ohmae, Javier F Medina, Reza Shadmehr

**Affiliations:** Laboratory for Computational Motor Control, Dept. of Biomedical Engineering, Johns Hopkins School of Medicine, Baltimore, Maryland; School of Cognitive Sciences, Institute for Research in Fundamental Sciences, Tehran, Iran; Department of Physiology and Biophysics, Washington National Primate Center, University of Washington, Seattle, Washington; Memory and Brain Research Center, Dept. of Neuroscience, Baylor College of Medicine, Houston, Texas; Department of Otolaryngology – Head and Neck Surgery, Washington National Primate Center, University of Washington, Seattle, Washington

## Abstract

Analysis of electrophysiological data from Purkinje cells (P-cells) of the cerebellum presents challenges for spike detection. Complex spikes have waveforms that vary significantly from one event to the next, raising the problem of misidentification. Even when complex spikes are detected correctly, the simple spikes may belong to a different P-cell, raising the danger of misattribution. Here, we analyzed data from over 300 P-cells in marmosets, macaques, and mice, using an open-source, semi-automated software called P-sort that addresses the spike identification and attribution problems. Like other sorting software, P-sort relies on nonlinear dimensionality reduction to cluster spikes. However, it also uses the statistical relationship between simple and complex spikes to merge seemingly disparate clusters, or split a single cluster. In comparison with expert manual curation, occasionally P-sort identified significantly more complex spikes, as well as prevented misattribution of clusters. Three existing automatic sorters performed less well, particularly for identification of complex spikes. To improve development of analysis tools for the cerebellum, we provide labeled data for 313 recording sessions, as well as statistical characteristics of waveforms and firing patterns.

## Introduction

Recording neuronal activity from the cerebellum presents both opportunities and challenges. The principal cells of the cerebellum, Purkinje cells (P-cells), can be identified based on their unique electrophysiological properties. Among cells in the cerebellum, only P-cells can produce simple and complex spikes (Thach, 1967). This makes it possible to use statistical methods to measure the likelihood that the recorded neuron is a P-cell: generation of a complex spike should be followed by suppression of simple spikes (Eccles et al., 1966; Sato et al., 1992). However, detection of complex spikes is difficult because these spikes are not only rare, but their waveforms also vary from one spike to the next. Thus, it is common to detect the simple spikes but not the complex spikes, or alternatively, detect the complex spikes but later realize that they are not followed with simple spike suppression and therefore do not belong to the same P-cell. To address these issues, we developed a spike analysis software that aids detection of simple and complex spikes, as well as quantifies whether the two events are generated by a single P-cell.

Unlike simple spikes, the power spectrum of complex spikes tends to be greatest in the low-frequency range (30-800 Hz). As a result, a typical complex spike can produce a “broad spike” in the low-pass filtered representation of the data (local field potential, LFP). Indeed, two recent developments in complex spike detection are novel algorithms that depend partly on the LFP waveform (Markanday et al., 2020; Zur & Joshua, 2019). Once the simple and complex spikes are labeled, the final step is to determine whether the simple spikes have been suppressed after a complex spike. If so, then one may conclude that the two kinds of spikes were generated by a single P-cell. However, in some data sets complex spikes do not have an LFP signature. Moreover, even if the complex spikes are detected, the detected simple spikes may belong to a different P-cell, or even a non P-cell.

As a result, the problem is two folds: in the identification step, we need to label the simple and complex spikes, whereas in the attribution step, we need to determine which group of complex spikes was generated by the P-cell that produced a particular cluster of simple spikes. To consider these challenges, we formed a collaboration that included laboratories which focused on marmosets, macaques, and mice. Our software was developed using a database of over 300 P-cells recorded in three species.

The diversity of species and recording electrodes helped us identify some of the critical issues that are present in cerebellar electrophysiology. The presence of experts from the various laboratories provided a diversity of opinions, helping us verify the algorithms, as well as highlight their limitations. Here we report the results of this effort.

P-sort is an open-source, Python-based software that runs on Windows, MacOS, and Linux platforms. To cluster waveforms and identify simple and complex spikes, P-sort uses both a linear dimensionality reduction algorithm and a novel nonlinear algorithm called UMAP (Uniform Manifold Approximation and Projection) (McInnes et al., 2018). Importantly, it quantifies the probabilistic interaction between complex and simple spikes, providing an objective measure that can split a single cluster, or merge two different clusters, despite similarities or differences in their waveforms. Thus, P-sort helps the user go beyond waveforms to improve clustering of spikes.

However, a limitation of P-sort is that it relies on user interaction. To encourage development of more automated algorithms for the cerebellum, with this report we provide a large database of labeled spikes from all three species. P-sort’s source code is available at:

https://github.com/esedaghatnejad/psort. The labeled data are available at: https://doi.org/10.17605/osf.io/gjdm4.

## Results

To illustrate the variety of challenges that we face in cerebellar neurophysiology, consider the data shown in Fig. 1. Here, the LFP channel (10-200 Hz) is plotted in red, and the AP channel (50-5000 Hz) is plotted in black. Occasionally, one is lucky enough to isolate a P-cell that exhibits easily identifiable complex and simple spikes, as shown in Fig. 1A. In this example, LFP shows a large positive peak for the complex spike. To confirm that the complex and simple spikes originate from the same cell, we compute the conditional probabilities Pr(S(t)|C(0)), and Pr(S(t)|S(0)), over a domain of ±50 ms. The term Pr(S(t) | C(0)) indicates the probability that a simple spike occurred at time *t*, given that a complex spike was generated at time zero. The term Pr(S(t) | S(0)) is the probability of a simple spike at time *t*, given that another simple spike was generated at time zero. Spikes that originate from a single cell produce a suppression period (Gao et al., 2012). Thus, Pr(S(t)|S(0)) exhibits a near zero probability period centered at time zero. On the other hand, a complex spike coincides with the suppression of future (but not past) simple spikes. As a result, Pr(S(t) | C(0)) is asymmetric, with a long period of near zero simple spike probability following the time point zero. The presence of simple spike suppression following a complex spike, as shown in Fig. 1A (right pannel), confirms that these two groups of spikes are generated by the same P-cell.

**Fig. 1.**
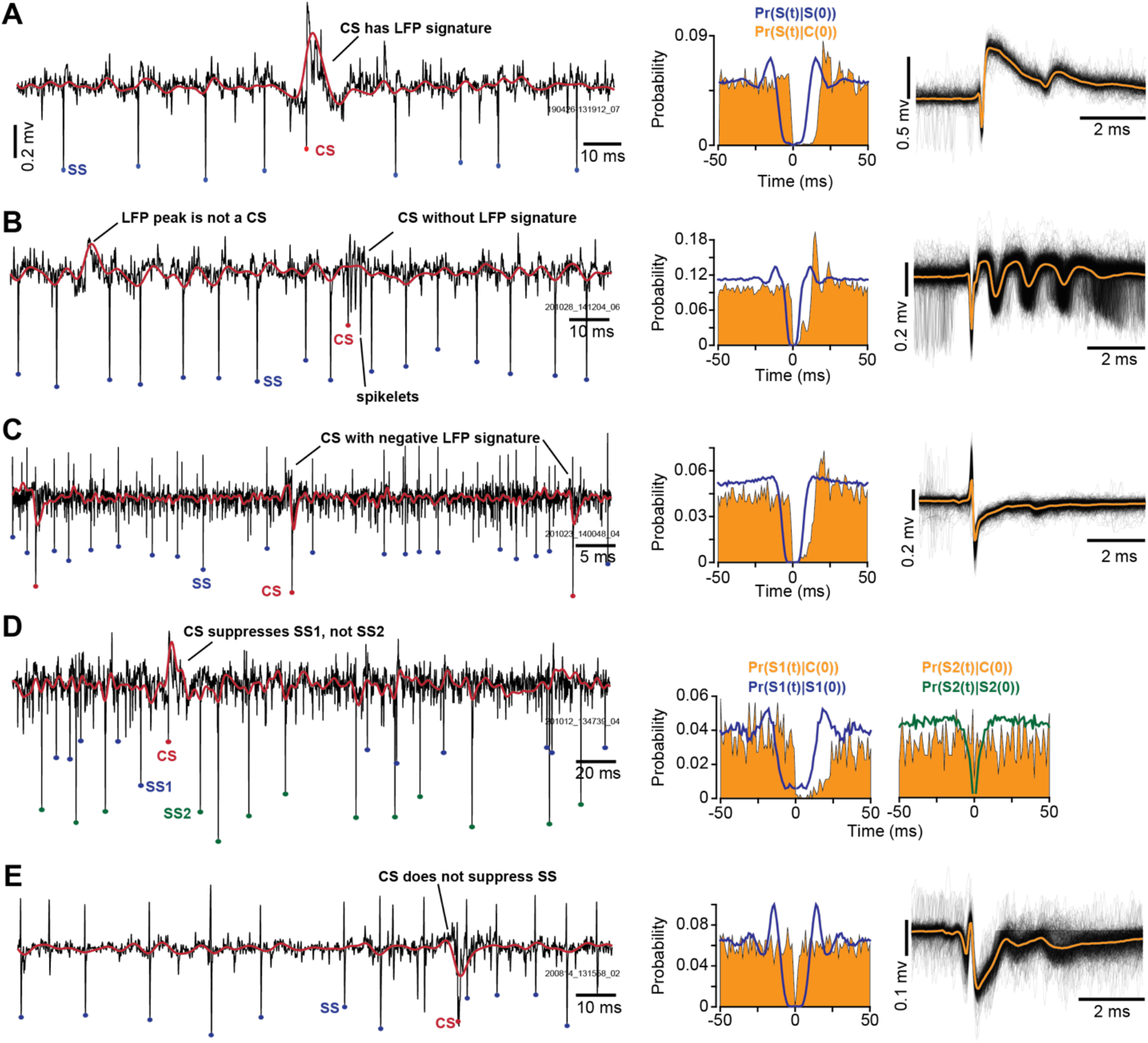
Examples of challenges in cerebellar spike sorting. In the left column, LFP channel (10-200 Hz) is plotted in red, and the AP channel (50-5000 Hz) is plotted in black. The middle column displays the conditional probability of a simple spike at time *t*, given that a complex spike occurred at time zero, labeled as Pr(S(t)| C(0). Note the asymmetric suppression following a complex spike. The conditional probability Pr(S(t) | S(0)) indicates the probability of a simple spike at time *t*, given that another simple spike occurred at time zero. The right column includes individual complex spike traces, as well as the average trace. **A**. In this recording, complex spikes have a positive LFP peak. **B**. Here, complex spikes cannot be identified from their LFP waveform as they lack an LFP signature. **C**. In this recording, complex spikes have a negative LFP peak. **D**. Here, complex spikes coincided with the suppression of one group of simple spikes (SS1), but not a second group (SS2). The probability pattern (right column) suggests that SS2 is likely not a P-cell. **E**. In this recording, complex spikes do not coincide with suppression of the simple spikes. Thus, the two groups of spikes are not generated by the same P-cell. Bin size is 1 ms for the probability plots.

Fig. 1B presents a more challenging example. Here, the complex spikes do not have an LFP signature, and thus are unlikely to be detected in the LFP channel. However, analysis of the AP channel using a novel nonlinear dimensionality reduction algorithm called UMAP (McInnes et al., 2018) identifies potential complex spikes. The identified events are genuine complex spikes, as evidence by their spikelets (more examples from the same cell are shown in Fig. 2A), and the fact that the simple spikes have been suppressed after a complex spike (Fig. 1B, right panel).

**Fig. 2.**
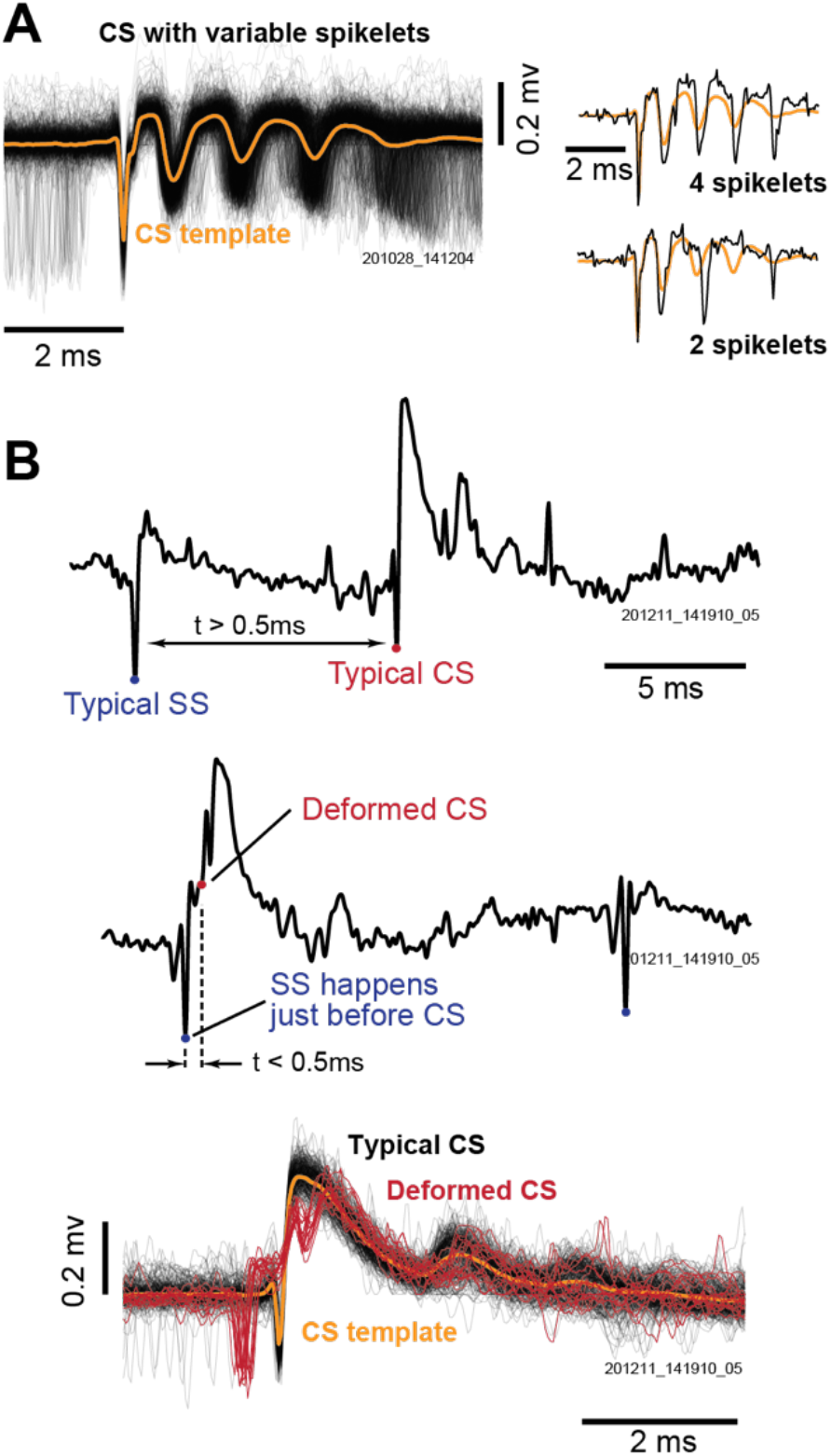
Additional challenges in identification of complex spikes. **A**. Complex spike waveforms vary because of the timing and number of spikelets. **B**. Complex spike waveforms vary because of the temporal proximity of a simple spike. Here, a simple spike before onset of a complex spike distorts the complex spike waveform. Top and middle rows: simple spike occurs at 3 or 0.2 ms before a complex spike. Bottom row: red traces are complex spikes with deformed waveforms.

In a well isolated P-cell, the complex spikes can produce spikelets that are similar to simple spikes. For example, the complex spike in Fig. 1B exhibits large spikelets, events that may be difficult to dissociate from ordinary simple spikes. This is evidenced by the fact that Pr(S(t) |C(0)) shows a small non-zero probability between 0 and +10 ms (right panel of Fig. 1B), during the period in which we would expect a near complete suppression.

Another example of the diversity of complex spike waveforms is shown in Fig. 1C. In this case, the complex spike exhibits a negative LFP peak. Nevertheless, once the complex spikes are correctly identified, the simple spikes followed a suppression period.

While detection of complex spikes may be challenging, a more crucial problem is attribution: sometimes the prominent group of simple spikes belongs to one P-cell, while the complex spikes belong to a different P-cell. An example of this problem is shown in Fig. 1D. Here, the LFP signal allows for detection of the complex spikes. However, there are two groups of simple spikes, SS1 and SS2. The spikes labeled SS2 are the larger amplitude events, but these spikes are not followed by a suppression period after the complex spikes. Rather, the smaller amplitude events SS1 are the spikes that followed by a suppression after the complex spikes.

A different form of the attribution problem is shown in Fig. 1E. In this case, the simple spikes are easily identified, and the complex spikes have a negative LFP. Remarkably, despite the excellent isolation, the simple spikes have not been suppressed after the complex spikes. Rather, the two groups of spikes in this recording are generated by two different P-cells.

Identification of complex spikes suffers from additional problems. There are variable number of spikelets in the complex spike waveform (Burroughs et al., 2017; Davie et al., 2008; Ito & Simpson, 1971; Monsivais et al., 2005; Najafi & Medina, 2013; Yang & Lisberger, 2014), and thus template matching may have difficulty labeling all complex spikes (Markanday et al., 2020). Examples of the variable spikelets are shown in Fig. 2A. Paradoxically, the better the P-cell’s isolation, the larger the impact of waveform variations caused by spikelets, and thus the greater the risk that some complex spikes will be missed.

Finally, the complex spike waveform can be distorted because of the proximity of a simple spike, as shown in Fig. 2B (Markanday et al., 2020; Servais et al., 2004). This is because simple and complex spikes are driven by different inputs to the P-cell, one is from a climbing fiber that generates a dendritic complex spike, while the other is from granule cells that generate somatic simple spikes. As a result, a simple spike can be generated up to a fraction of a millisecond before a complex spike (middle plot of Fig. 2B). Here, the complex spike that follows the simple spike at short latency lacks the sharp component that initiated more typical complex spikes. As a result, the complex spike waveform is distorted by the proximity of the simple spike (lower plot, Fig. 2B).

In summary, identification of complex spikes may be difficult because they can lack an LFP signature, their waveforms can be distorted because of nearby simple spikes, or their waveforms can incorporate variable number of spikelets. After the simple and complex spikes are identified, one still faces the problem of finding the simple spike cluster that belongs to the P-cell that generated the complex spikes. We built P-sort to help with these identification and attribution problems.

### Clustering waveforms

The diversity of complex spike waveforms suggests that it may be difficult to find a single mathematical technique that could identify these spikes in all situations. For example, in some cases it is possible to identify the complex spikes from the LFP channel (Fig. 1A), whereas in other cases it is necessary to search the AP channel (Fig. 1B). In the case where two types of simple spikes are present, often the larger amplitude simple spikes get suppressed after the complex spikes (Fig. 1C), but occasionally the smaller amplitude spikes are the correct choice (Fig. 1D). P-sort provides tools for clustering as well as hypothesis testing. The tools include traditional dimensionality reduction methods such as principal component analysis (PCA), as well as novel algorithms such as UMAP (McInnes et al., 2018). As the user identifies putative groups of simple and complex spikes, the software provides immediate statistical feedback regarding the probability that the spikes are from the same P-cell. Thus, the main idea of P-sort is to merge the identification and attribution steps into a single framework.

P-sort works best with the raw, broad-spectrum recording such as the data generated by widely used Open Ephys, an open-source data acquisition software (Siegle et al., 2017). However, P-sort also works with data in which the LFP and AP channels are acquired separately. Upon loading the data, the user is provided with a GUI to specify how the data should be chunked into “slots”. Each slot is a region of data that is analyzed in turn, but once a slot is analyzed, other subsequent slots inherit features such as spike templates. In case of broad-spectrum data, the user can specify the filter properties for the LFP and AP channels (Fig. 3A, part 1). Once these filters are selected, P-sort automatically selects a threshold for each channel (Fig. 3, part 2, see Methods) and displays the statistics of the resulting simple and complex spikes (Fig. 3, parts 3 and 4).

**Fig. 3.**
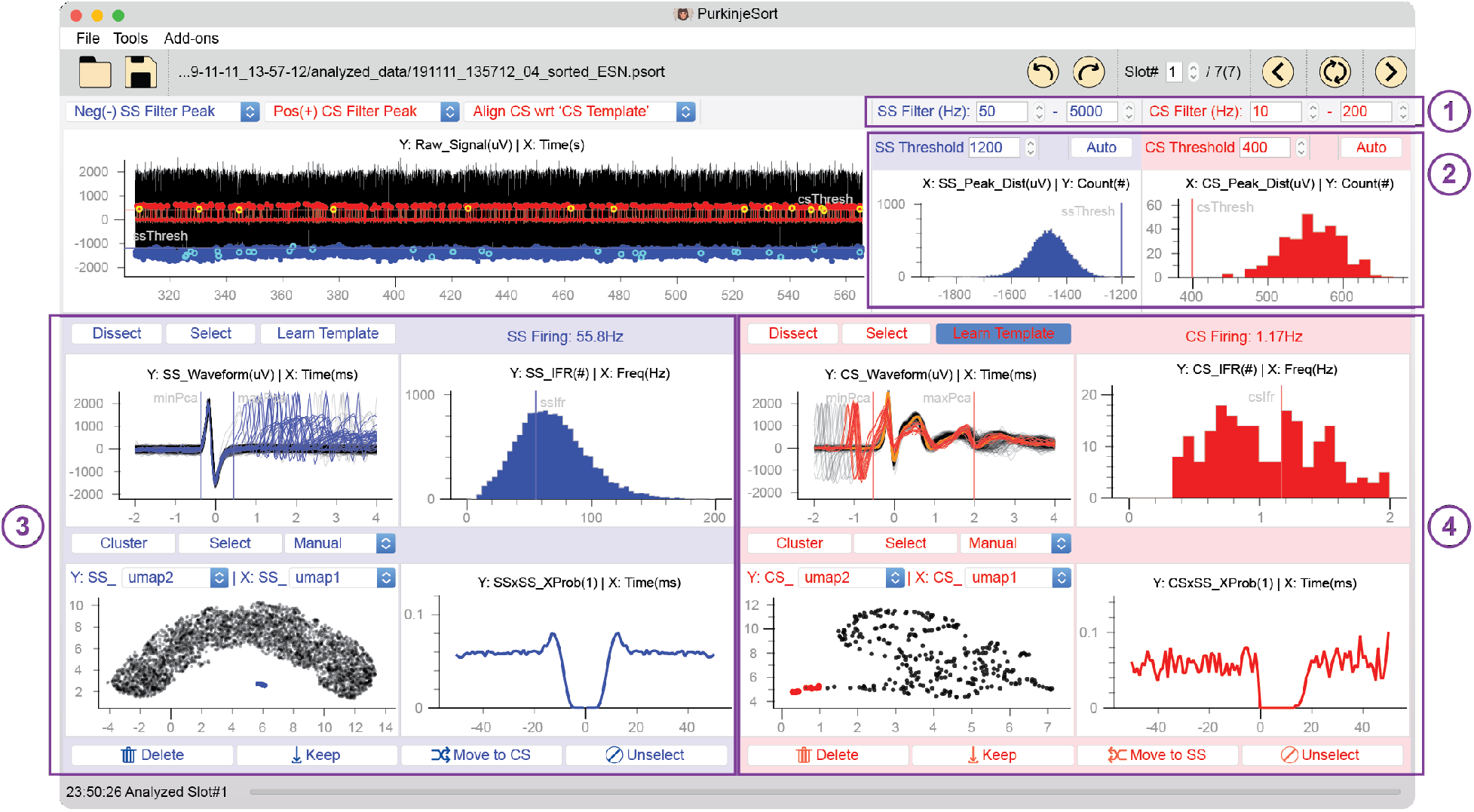
Main window of P-sort. **1**. Specification of filter properties for the AP and LFP channels. **2**. Threshold selection and the distribution of voltage peaks for events detected in the LFP and AP channels. **3**. Identification of simple spikes. The windows include spike waveforms aligned to peak voltage, distribution of instantaneous firing frequency, conditional probability Pr(S(t)|S(0)) demonstrating a suppression period, and clustering using UMAP. **4**. Identification of complex spikes. The windows include spike waveforms aligned to a learned template (yellow), distribution of instantaneous firing frequency, conditional probability Pr(S(t) | C(0)) demonstrating suppression of simple spikes, and clustering using UMAP. The red traces indicate complex spikes that have been distorted because of nearby simple spikes.

In the example shown in Fig. 3, P-sort automatically selected thresholds for LFP and AP channels and identified simple and complex spike candidates. The UMAP space indicated a single simple spike cluster (Fig. 3, part 3), which was confirmed by Pr(S(t) | S(0)), illustrating that the simple spikes had around 10 ms suppression period (centered at zero). However, the UMAP space was not homogeneous for the complex spikes (Fig. 3, part 4), meaning that there was variability in the waveforms. Regardless, Pr(S(t) | C(0)) exhibited around 15 ms suppression period. Thus, there was statistical confirmation that the simple spikes were well isolated, and they have been suppressed after the complex spikes.

Of course, in most cases the data are not as easily sorted as the case shown in Fig. 3. Another frequent case is one in which the complex spikes do not have an LFP signature (Fig. 4A), and thus one must focus the search on the AP channel. However, in this recording the simple and complex spike waveforms happen to be quite similar (Fig. 4C). As a result, in the PCA space the data present a single cluster (Fig. 4B, left subplot). Thus, if we were to rely on PCA alone, we might conclude that only simple spikes are present.

**Fig. 4.**
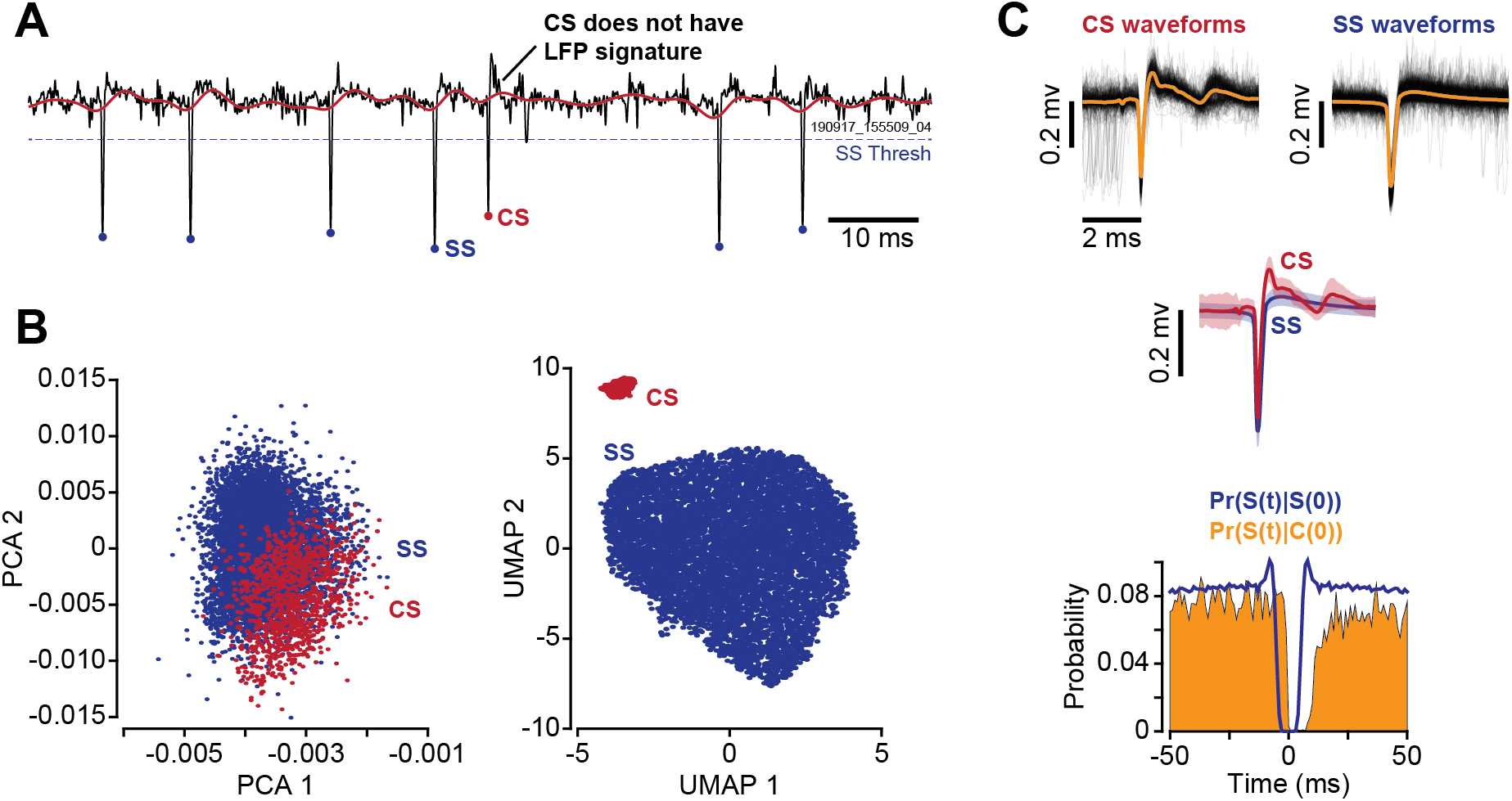
UMAP dissociates simple and complex spikes. **A**. In this recording, complex spikes do not exhibit an LFP signature (red trace). **B**. Clustering of the spikes in PCA space does not produce a clear separation. However, the two groups of spikes separate in the UMAP space. The complex spikes identified by UMAP are shown in red in the PCA space. **C**. The waveforms and average traces for the complex spike and simple spike clusters as identified by UMAP. The conditional probabilities demonstrate that the complex spike events coincide with suppression of the simple spikes, suggesting that the two groups of spikes are likely generated by the same P-cell. Error bars are standard deviation.

P-sort utilizes a novel dimensionality reduction algorithm called UMAP (McInnes et al., 2018). UMAP is a nonlinear technique that, in our experience, is particularly powerful for clustering waveforms and identifying complex spikes, as also shown by the work of Markanday et al. (2020). Indeed, in the case of the data in Fig. 4, projecting the waveforms onto the UMAP space unmasks two clusters (Fig. 4B, right subplot). Using the graphical user interface (GUI) we select the smaller group of spikes and tentatively label them as complex spikes. Immediately, P-sort updates the probability Pr(S(t)|C(0)) window, as shown in Fig. 4C, illustrating that these putative complex spikes were followed by simple spike suppression.

One of the issues in identifying complex spikes is that the waveform can be significantly distorted by a preceding simple spike. Indeed, a P-cell can produce a complex spike at a fraction of a millisecond following a simple spike, as shown in Fig. 2B. The simple spike proximity distorts the complex spike waveform, making it difficult to correctly locate onset of the complex spikes. If the distortion is small, template matching can still identify the onset of the complex spikes (Fig. 3, parts 4). However, as we will see below, template matching can sometimes produce an incorrect alignment of complex spikes.

An example of this situation is shown in Fig. 5A. Initially, the putative complex spikes are aligned by P-sort based on a sodium peak which resembles simple spikes (see Methods). This alignment results in two groups of complex spikes (shown by black and red traces in Fig. 5A). Next, the user defines a new template based on the mean waveform of the correctly aligned complex spikes (Fig. 5A, black traces) and resort the data (Fig. 5B). Despite significant improvement in the performance of the complex spike alignment, the problem of aligning the deformed complex spikes still persists. Indeed, the UMAP space indicates presence of two groups of complex spikes (red and black dots in Fig. 5B). However, because both complex spike groups coincided with simple spike suppression, they are likely a single cluster that need to be merged. To correct the error, P-sort provides the Dissect Module. The Dissect Module is a semi-manual platform to re-label the onset of the distorted complex spikes (Fig. 5C), and correctly identify the simple spikes that precede them. Following this re-labeling, the complex spikes are correctly aligned (Fig. 5D).

**Fig. 5.**
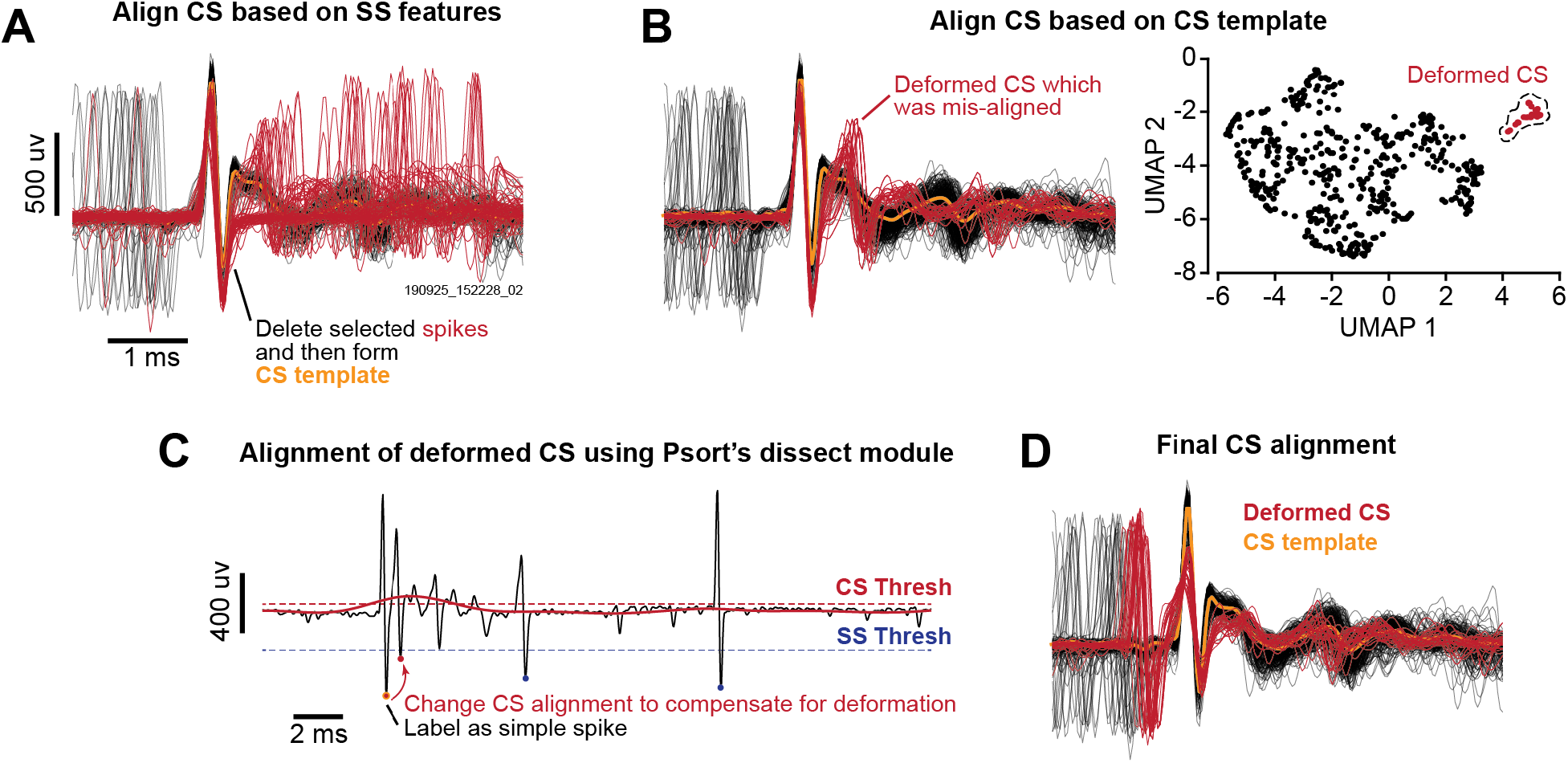
Correcting for misalignment of complex spikes. **A**. P-sort initially aligns complex spikes based on the sodium/potassium peak which resembles simple spikes (see Methods). This results in misalignment of some complex spikes because these complex spikes do not express sodium/potassium peak. The user can delete the misaligned spikes (red) and compute a complex spike template (yellow). **B**. Alignment of the spikes to the complex spike template does not solve the problem for all complex spikes: some of the deformed complex spikes remain misaligned. **C**. P-sort provides a Dissect Module with which the user can change the spike alignment to compensate for the deformation. **D**. Final complex spike alignment. Note the deformation in the complex spikes caused by the proximity of the simple spikes (red traces).

### Merging clusters or splitting them using statistical interactions between simple and complex spikes

A unique challenge in cerebellar neurophysiology is finding the simple and complex spike clusters that belong to a single P-cell. P-sort provides cluster labeling and statistical tools to help with this problem. However, the main strength of P-sort is to go beyond waveform clustering and use the statistical relationship between complex and simple spikes to identify an additional feature that can help merge clusters, or even split a single cluster.

To illustrate this, let us begin with a particularly challenging example as shown in Fig. 6A. Here, a visual inspection of the waveforms suggests presence of multiple neurons. Indeed, UMAP clustering produces numerous groups of simple (Fig. 6B) and complex spikes (Fig. 6C). The task is to find the clusters that are signal and not noise, and more importantly, determine which simple spike cluster(s) can be attributed to which complex spike cluster(s).

**Fig. 6.**
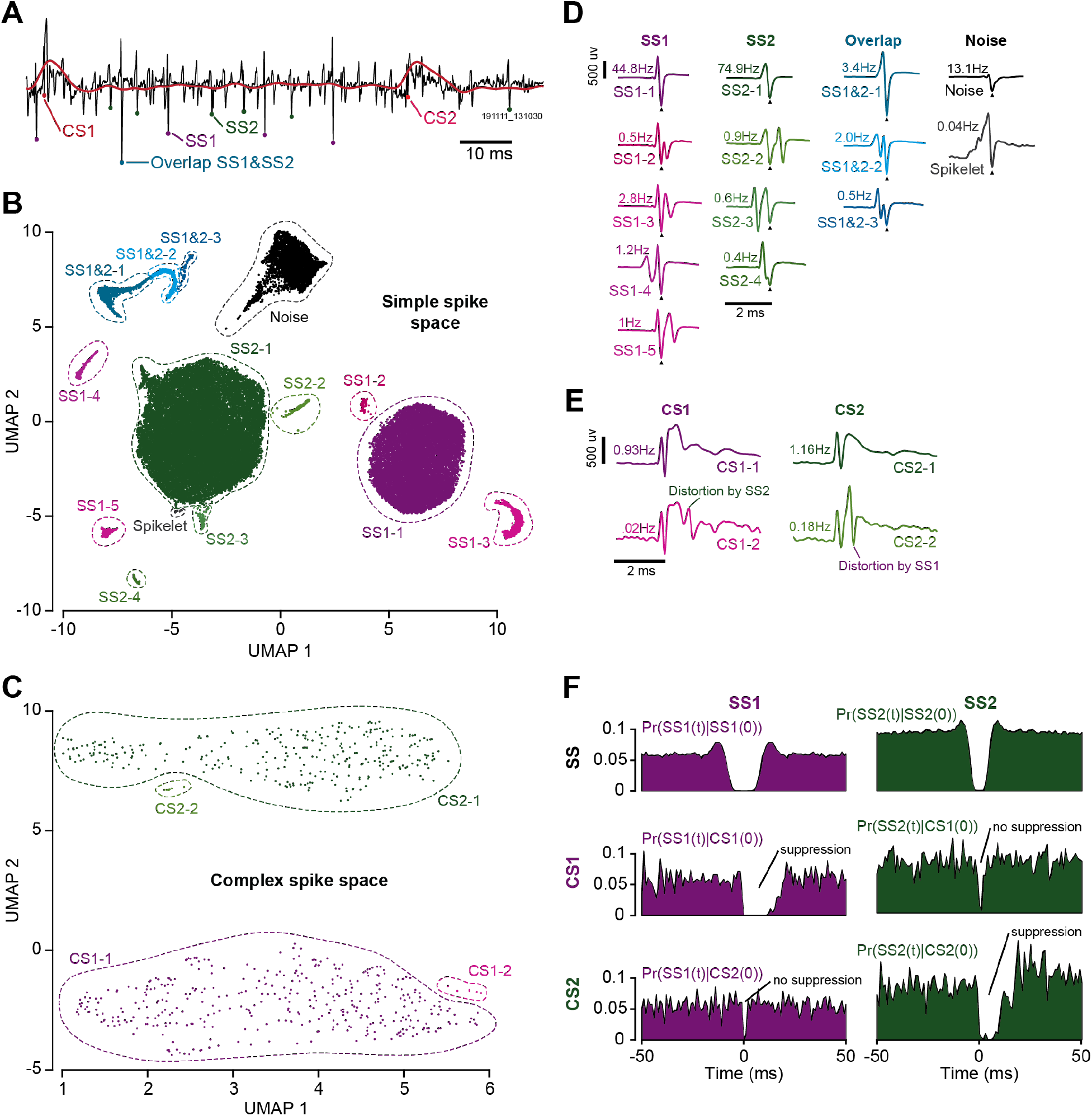
Finding clusters of simple and complex spikes that are likely generated by the same P-cell. **A**. This recording includes at least four groups of spikes: two that appear to be complex spikes, and two that are simple spikes. **B**. UMAP clustering of the simple spike space. The two major clusters are SS1 and SS2. Their waveforms are shown by SS1-1 and SS2-1 in part D. The smaller clusters are distorted spikes that are due to the temporal proximity of these major spikes, as well as other spikes, as shown in part D. A smaller cluster of spikes are labeled as spikelets of complex spikes. **C**. UMAP clustering of the complex spike space. The two major clusters are CS1 and CS2. Their waveforms are labeled as CS1-1 and CS2-1 in part E. Their waveforms can be distorted by simple spikes, as shown by CS1-2 and CS2-2. **D**. Waveforms of various clusters labeled in the simple spike space. **E**. Waveforms of the four clusters labeled in the complex spike space. **F**. Suppression period of SS1 and SS2 is quantified by the probability Pr(SS1(t) | SS1(0)) and Pr(SS2(t) | SS2(0)). The probability Pr(SS1(t) | CS1(0)) quantifies the suppression following CS1. Thus, CS1 coincides with suppression of SS1 but not SS2. CS2 coincides with suppression of SS2 but not SS1. Bin size is 1 ms.

The reason for the numerous clusters in the simple spike UMAP space (Fig. 6B) is because the electrode is picking up signals from multiple neurons, and sometimes spikes from one neuron can distort the spike from another neuron. For example, the cluster labeled SS1-1 in Fig. 6B is due to simple spikes from neuron 1 (Fig. 6D, SS1-1). The nearby cluster SS1-2 in Fig. 6B is due to simple spikes from neuron 1 that are in close temporal proximity with a spike from another neuron (Fig. 6D, SS1-2). There are also clusters associated with spikes from neuron 2 that occur in isolation (Fig. 6D, SS2-1), or in close temporal proximity with a spike from another neuron (Fig. 6D, SS2-2). Sometimes, SS1 and SS2 co-occur, resulting in a larger spike, as labeled by cluster SS1&2-1. Finally, there are spikelets in the complex spike waveform that can be mis-labeled as simple spikes. The spikelets are noted in Fig. 6D.

There are also four clusters in the complex spike UMAP space (Fig. 6C). One large cluster is associated with complex spikes labeled CS1 (Fig. 6E, CS1-1), while the other large cluster is associated with a second group of complex spikes labeled CS2 (Fig. 6E, CS2-1). Near each of these large clusters there are smaller clusters, reflecting the variability in the complex spike waveform. A source of variability in the complex spike waveform is presence of simple spikes from multiple P-cells. Thus, the complex spike will coincide with suppression of one group of simple spikes, but not all groups. As a result, CS1 can be distorted by arrival of simple spike labeled SS2, and CS2 can be distorted by arrival of simple spikes labeled SS1 (Fig. 6E, CS1-2 and CS2-1).

P-sort provides tools to label the clusters, examine their statistical properties, and determine whether clusters should be merged or not. For example, both the conditional probabilities and the statistics of the firing rates suggest that SS1 and SS2 are two different simple spikes, as shown in Fig. 6F, first row. Furthermore, the complex spike cluster CS1 coincided with the suppression of simple spike cluster SS1, but not SS2 (Fig. 6F, second row). Similarly, the complex spike cluster CS2 coincided with the suppression of simple spike cluster SS2, but not SS1 (Fig. 6F, third row). As a result, in this recording we have two distinct P-cells.

In a second example, let us show that the statistical interactions between simple and complex spikes can provide evidence suggesting that a single cluster of spikes may in fact be composed of two different cells. This data set is shown in Fig. 7. In this recording, the complex spike waveforms produce a single cluster in the UMAP space (Fig. 7A). However, P-sort notes that only a part of this complex spike cluster coincided with the suppression of the simple spikes. The sub-cluster that proceeded the suppression of the simple spikes is labeled as CS1, and its waveform is shown in Fig. 7B. Notably, the sub-cluster CS2 has a waveform that is similar to CS1, but CS2 does not coincided with the suppression of the simple spikes, as illustrated in Fig. 7C.

**Fig. 7.**
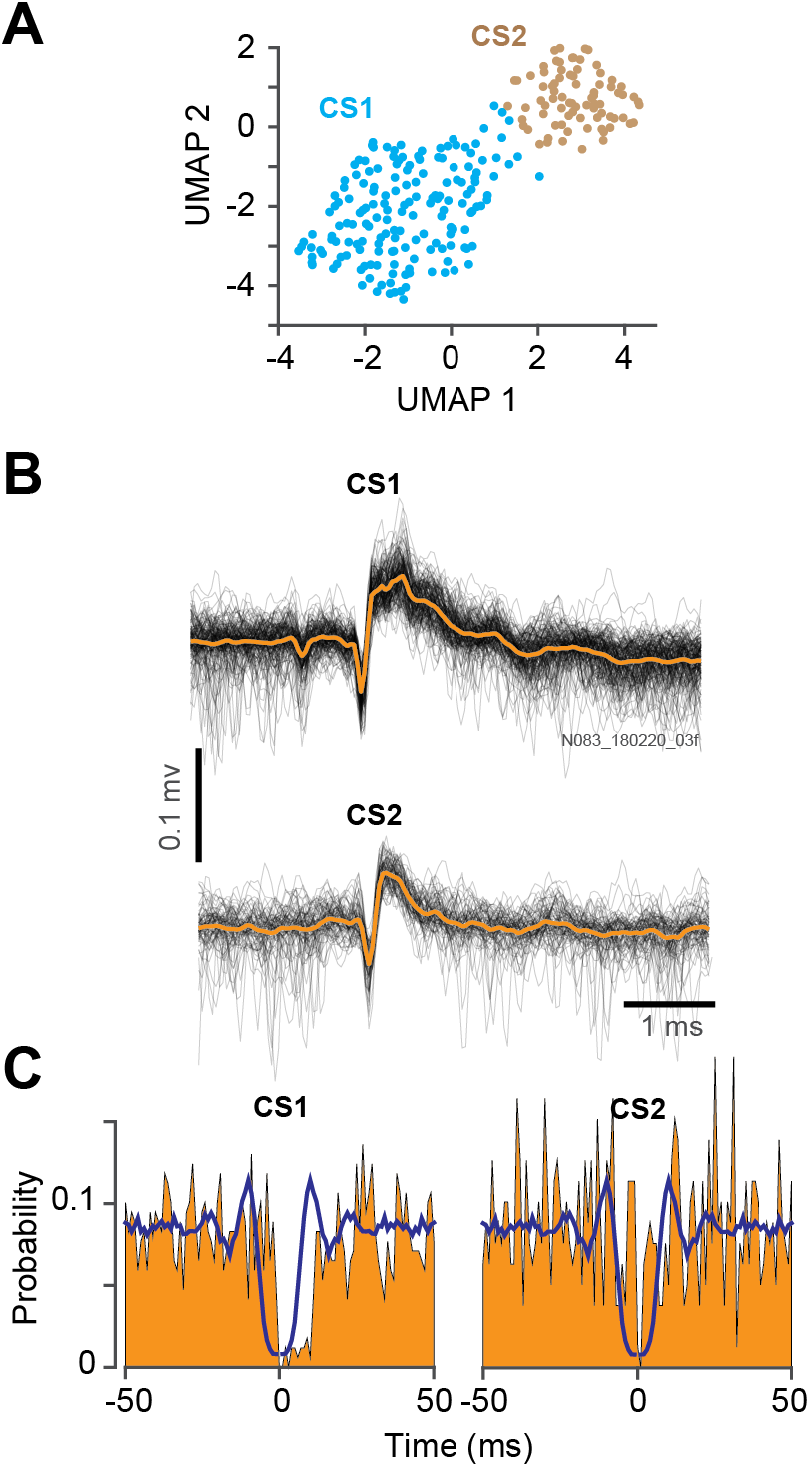
Splitting a complex spike cluster based on its statistical properties with simple spikes. **A**. In this dataset, the UMAP space indicates a single complex spike cluster. However, statistical considerations raise the possibility of two subgroups, CS1 and CS2. **B**. The waveforms for the CS1 and CS2 subgroups are similar. **C**. The yellow regions are the conditional probabilities of simple spike suppression following a CS1 at time zero, or CS2 at time zero. Note that CS1 coincides with suppression of the simple spikes, but CS2 does not. Thus, while it is difficult to split the complex spike cluster into two groups based on their waveforms, they exhibit very different statistical properties.

This example highlights the possibility that on occasion, the complex spike waveforms may suggest presence of a single cluster, but a consideration of the statistical interactions can reveal that the simple spikes have been suppressed by only a sub-group within that cluster. This interaction between simple and complex spikes is a crucial feature that is utilized by P-sort to go beyond waveforms to help correctly identify and attribute spikes of a P-cell.

In summary, P-sort provides clustering tools to identify simple and complex spikes. It relies on template matching to find onset of spikes, but also provides tools to correct instances where template matching can fail. A critical feature of P-sort is to provide tools for labeling and pairing of simple and complex spikes, thus allowing the user to visualize the likelihood that specific groups of spikes are generated by a single P-cell. This can lead to merging of nearby complex spike clusters because both coincided with the suppression of the simple spikes, or splitting of a single cluster because only a sub-group coincided with the suppression of the simple spikes.

### Comparison of P-sort with expert manual curation

A common method currently employed for sorting of cerebellar data is via manual curation by an expert user, for example via Spike2. To measure the quality of P-sort results, we analyzed data from marmosets, mice, and macaques, and then compared the P-sort results with those generated by the experts in each laboratory.

Fig. 8A presents results from an example data set from the macaque cerebellum. In this data set, the expert and P-sort agreed on 25444 simple spikes (98.52% and 99.71%), and 555 complex spikes (98.58% and 83.33%). The median difference between the two methods in determining the timing of the spikes was 0.06±0.01 ms (median±MAD) for simple spikes, and −0.12±0.05 ms for complex spikes. P-sort labeled 74 (0.29%) simple spikes that were not identified by the expert. These are labeled as P-sort exclusive simple spikes. The expert labeled 382 (1.48%) simple spikes that were not identified by P-sort. These are labeled as expert exclusive simple spikes. P-sort’s rate of simple spikes was 45.5 Hz, while that of the expert was 46.0 Hz.

**Fig. 8.**
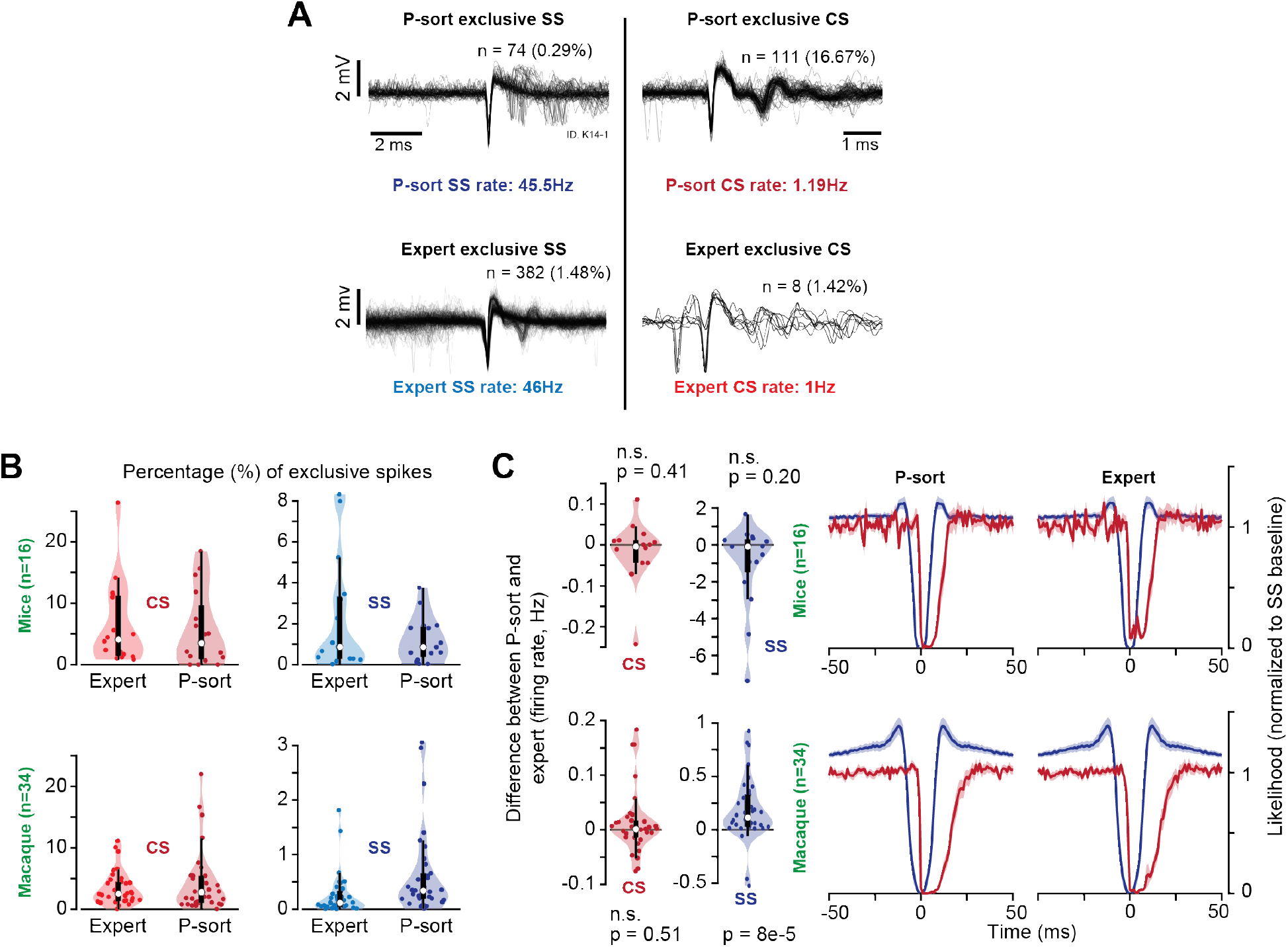
Comparison of P-sort with expert curation on mice and macaque data sets. **A**. Data from a macaque recording session. P-sort picked out 74 simple spikes that were not identified by the expert (0.29% of total), labeled as P-sort exclusive. Expert picked 382 simple spikes that were not identified by P-sort (1.48% of total), labeled as expert exclusive. Complex spikes that were exclusive to P-sort and the expert are also plotted. **B**. Summary statistics on the mice (n=16 sessions) and macaque (n=34 sessions) data sets. Percentage of exclusive simple and complex spikes are plotted for the expert and P-sort. The central mark indicates median of the distribution, and the bottom and top edges of the box indicate the 25^th^ and 75^th^ percentiles. The thin line indicates the range of the data excluding the outliers. **C**. Difference between P-sort and expert in terms of firing rate. Right columns show the likelihoods, normalized to the baseline simple spike probability in each session, averaged over all recording sessions for each species (bin size is 1 ms). Error bars are SEM.

For complex spikes, there were 111 (16.7%) events picked by P-sort that were not picked by the expert, producing a complex spikes rate of 1.19 Hz for P-sort vs. 1.0 Hz for the expert. Thus, P-sort identified 19% more complex spikes than the expert. The waveforms suggest that the P-sort exclusive complex spikes are likely valid. However, the 8 complex spikes picked by the expert and not P-sort are also likely valid, as indicated by their waveforms. The reason why P-sort missed these complex spikes is because in some cases, a simple spike was in temporal proximity and distorted the complex spike waveform. Thus, in this data set there was general agreement between the two methods.

In the macaque data set (34 sessions), a median of 2.53% of the complex spikes were detected exclusively by the expert, and 2.79% of the complex spikes were detected exclusively by P-sort. For simple spikes, 0.12% (median) of the spikes were detected only by the expert, as compared to 0.35% for P-sort. The two methods converged in their estimates of complex spike firing rates, but P-sort identified slightly more simple spikes (Fig. 8C, firing rate of P-sort spikes minus expert, Wilcoxon signed rank test, SS: Z=3.94, p=8e-5, CS: Z=0.65, p=0.51). The resulting conditional probabilities of the data sorted by P-sort and the expert were indistinguishable. Some of the complex spikes missed by P-sort were due to significant distortions that were present in the waveform. In general, agreement between P-sort and expert were higher for better isolated recording data.

In the mice data set (16 sessions), a median of 4.10% of the complex spikes were detected exclusively by the expert, and 3.53% of the complex spikes were detected exclusively by P-sort, as shown in Fig. 8B. For simple spikes, the agreement between the two methods was better: on average 0.86% (median) of the simple spikes were detected only by the expert, as compared to 0.857% for P-sort. Overall, there were no significant differences between P-sort and the expert in terms of rates of simple and complex spikes (Fig. 8C, firing rate of P-sort spikes minus expert, Wilcoxon signed rank test, SS: Z=-1.29, p=0.20, CS: Z=-0.82, p=0.41).

Importantly, in one recording session P-sort highlighted the possibility that the expert paired the wrong sub-group of complex spikes with the simple spikes. In this data set (Fig. 7), the expert labeled a single complex spike cluster, which is of course reasonable because of the similarity of the waveforms. However, P-sort split this cluster into two, labeling them as CS1 and CS2, and only attributed the CS1 sub-group as the complex spikes that were generated by the P-cell that also produced the simple spikes.

Overall, a comparison of P-sort with expert manual curation suggested a general agreement: the rates of simple and complex spikes were generally similar. However, for a few recording sessions P-sort was able to identify more complex spikes (Fig. 8A), and correctly label spikelets. In one instance, P-sort prevented pairing of the wrong subgroup of complex spikes with the simple spikes (Fig. 7).

### Comparison of P-sort with automated spike sorting

A major limitation of P-sort is that it is not automated, thus relying on the user to select tools and explore their efficiency in identifying and attributing spikes. In comparison, automated software can identify spikes with little or no user intervention. We compared P-sort’s results with three automated algorithms. In this comparison we included three data sets, one that was relatively easy, one that had medium difficulty, and one that was very difficult. Because the automated algorithms did not attribute pairs of simple and complex spikes, we manually performed this step following the conclusion of the automated spike sorting.

The simple and complex spikes for the easy and medium difficulty data sets are shown in Fig. 9A and Fig. 9B. The two kinds of spikes were easily separable in the first data set (Fig. 9A), but harder to separate in the second data set (Fig. 9B).

**Fig. 9.**
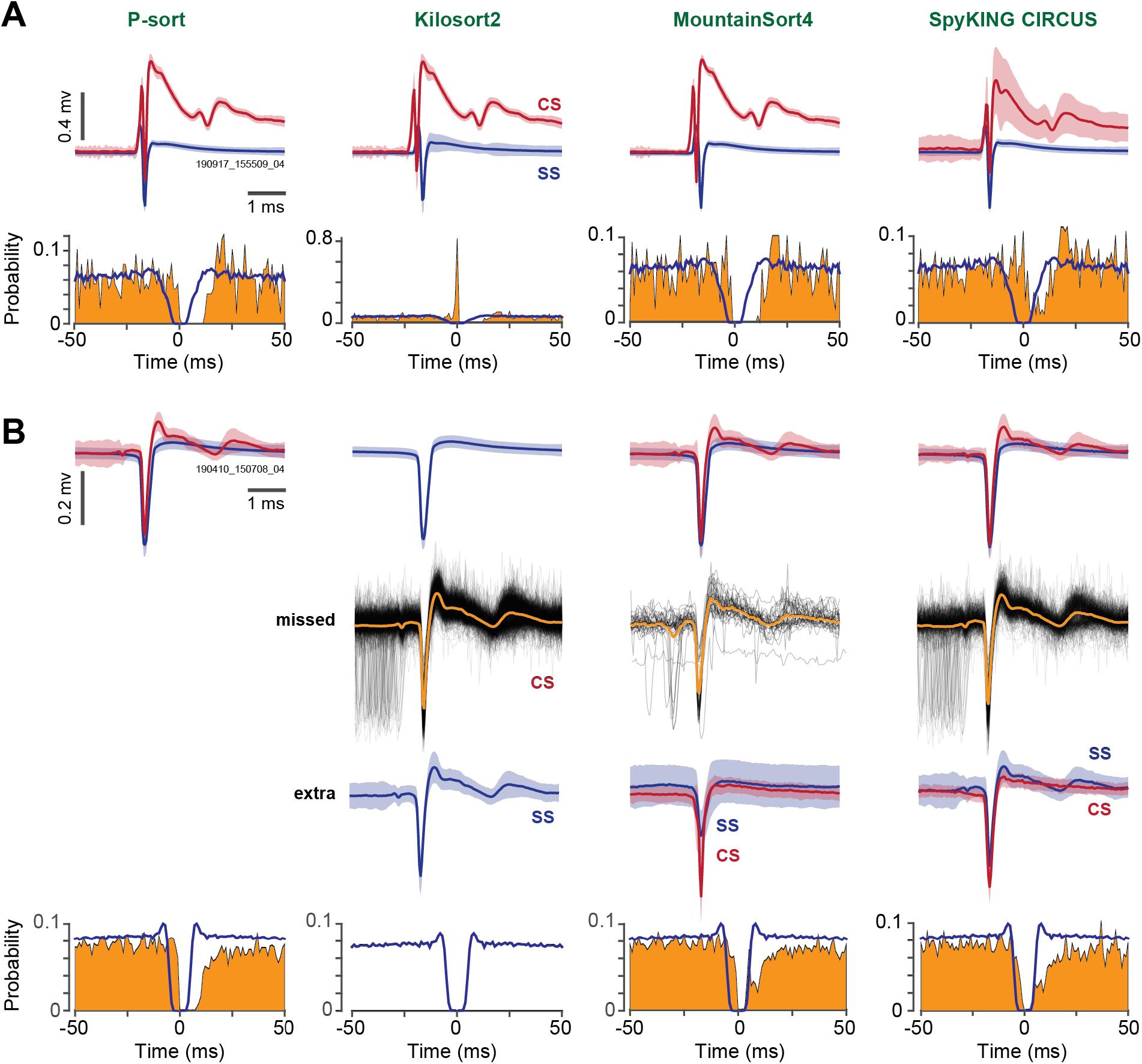
Comparison of P-sort with automated spike sorting algorithms on two data sets. **A**. Easy data set. The simple and complex spike waveforms are illustrated in the first row. The conditional probability for simple spikes Pr(S(t)|S(0)) is plotted in blue. The conditional probability for simple spike suppression following a complex spike Pr(S(t) | C(0)) is plotted in yellow. All algorithms identified the simple and complex spikes. Kilosort mislabeled the onset of the complex spike as a simple spike. **B**. More difficult data set. First row shows the spikes identified by each algorithm. Kilosort did not identify the complex spikes. Second row shows the complex spikes missed by each algorithm, with respect to P-sort. Third row shows the spikes that were identified by each algorithm but not P-sort. Fourth row is the conditional probabilities for the labeled spikes. Error bars are standard deviation.

In the easy data set, Kilosort2 (Pachitariu et al., 2016) found 10452 simple spikes and 146 complex spikes (first row of Fig. 9A), but mis-labeled a part of the complex spike waveform as a simple spike. This led to the unusual probability distribution shown in the second row of Fig. 9A. For the medium difficulty data set, Kilosort2 found the simple spikes, but did not identify any complex spikes (first row of Fig. 9B), resulting in missed spikes shown in the second row of Fig. 9B. It also mis-labeled many complex spikes as simple spikes (third row of Fig. 9B).

MountainSort4 (Chung et al., 2017) found 10460 simple spikes and 127 complex spikes in the easy data set, which compared well with 10461 simple spikes and 147 complex spikes identified by P-sort. In the medium difficulty data set, this automated algorithm found 70328 simple spikes and 996 complex spikes. For the simple spikes, the labeling was essentially identical with P-sort, but for the complex spikes the labeling differed by 178 events (17.2%). There were 36 P-sort identified complex spikes that were not labeled by MountainSort4 (second row of Fig. 9B), and some simple spikes that were labeled as complex spikes (third row of Fig. 9B). The labeling produced by P-sort resulted in a cleaner suppression period (Fig. 9B, bottom row), suggesting that in the medium difficulty data set, MountainSort4 mislabeled or missed around 15% of the complex spikes. The ability of MountainSort4 to find the complex spikes, but not Kilosort2, may be because MountainSort4 uses a PCA branching algorithm that is better in discrimination of local differences between waveforms in comparison to PCA.

SpyKING CIRCUS (Yger et al., 2018) found 10488 simple spikes and 198 complex spikes in the easy data set (Fig. 9A). Some (43/198) of the complex spikes found by SpyKING were probably not valid, as suggested by the relatively poor suppression period of simple spike illustrated by P(S(t) | C(0). In the medium difficulty data set, SpyKING found 70381 simple spike and 796 complex spikes, missing 493 complex spikes (second row of Fig. 9B) that were found by P-sort, and mislabeled some complex spikes as simple spikes (third row of Fig. 9B).

We also tested the algorithms on a very difficult data set that had multiple simple and complex spikes on a single contact, labeled as SS1, SS2, CS1, and CS2 in Fig. 6. Sorting this type of data can benefit significantly from the information in the statistical interactions between spike clusters. MountainSort4 found 48478 simple spikes SS1, and 51471 simple spikes SS2 (Supplementary Fig. 1A). This agreed with 98.6% of SS1 spikes labeled by P-sort, but only 67.4% of SS2 spikes. MountainSort4 found 1156 complex spikes CS1 and 892 complex spikes CS2. This agreed with 88.9% of CS1 and 85.3% of CS2 spikes labeled by P-sort. The complex spikes that were exclusively labeled by MountainSort4 or P-sort are shown on the right panel of Supplementary Fig. 1A. Some MountainSort4 CS1 spikes were labeled as CS2 by P-sort. Similarly, some MountainSort4 CS2 spikes were labeled as CS1 by P-sort. The conditional probabilities (Fig. 10) provide a method to compare these results. For SS1 spikes, the probability Pr(S(t)|S(0)) for P-sort (blue lines, Fig. 10, first row) exhibited a cleaner suppression period than MountainSort4. For the complex spikes, the probability Pr(S(t)|C(0)) for P-sort for SS1 by CS1, and SS2 by CS2, both showed a cleaner suppression for P-sort. Thus, the main disagreements in the two approaches were regarding the smaller amplitude simple spikes SS2, and memberships of complex spikes in CS1 and CS2.

**Fig. 10.**
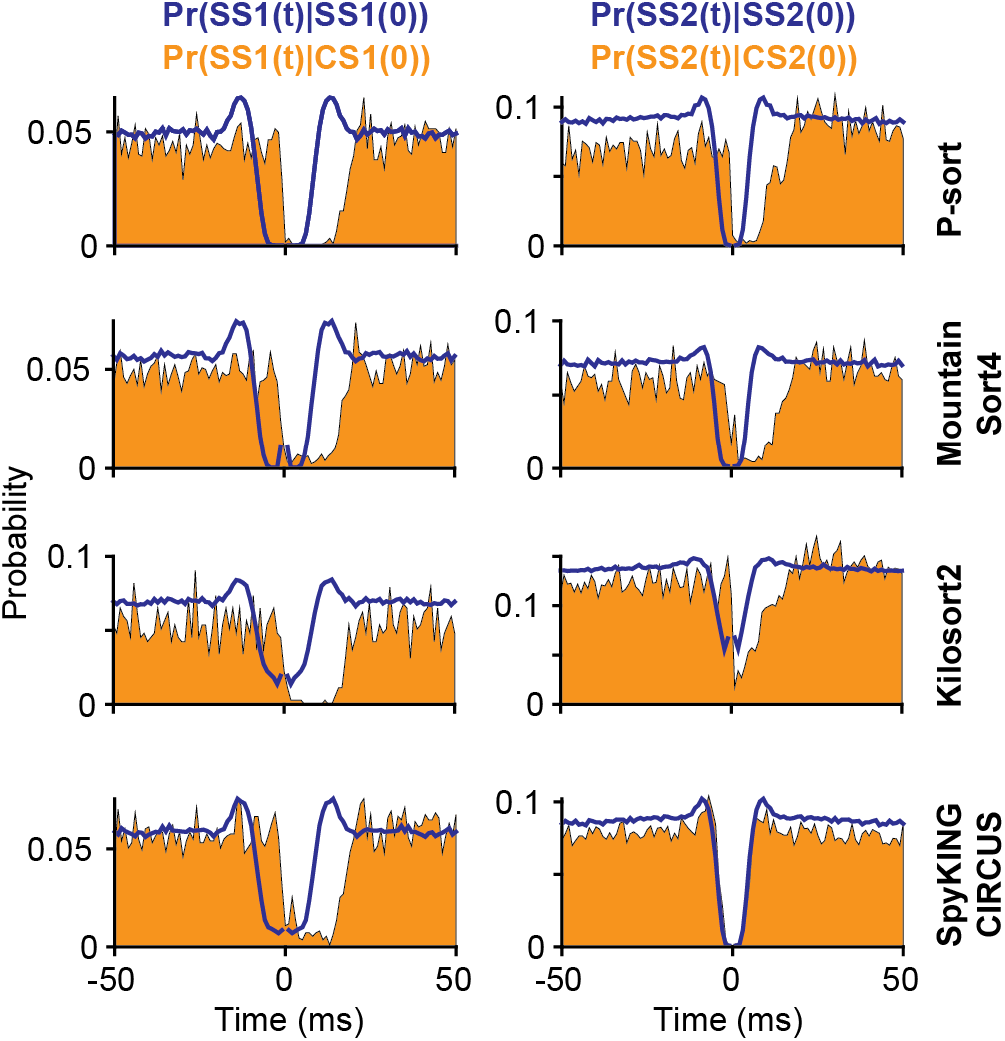
Performance of the automated algorithms on a difficult data set (Figs. 6). The plots show conditional probabilities for the simple and complex spikes identified by the automated algorithms and P-sort for the labels summarized in Fig. 10. Left column is the SS1 and CS1 relationship. Right column is the SS2 and CS2 relationship. For simple spikes, the suppression period is particularly poor for Kilosort2 for both SS1 and SS2. For complex spikes, SpyKING CIRCUS and Kilosort2 produce little or no suppression of simple spikes SS2. Bin size is 1 ms.

Kilosort2 found 56538 simple spikes SS1, and 117750 simple spikes SS2 (Supplementary Fig. 1B). The smaller magnitude spikes SS2 formed a much larger group in Kilosort2 as compared to both MountainSort4 and P-sort. For complex spikes, Kilosort2 found only 354 CS1, missing roughly 70% of the CS1 complex spikes labeled by P-sort. In contrast, it found 1117 CS2 events, agreeing with 94.3% of CS2 events found by P-sort. Many of the CS2 complex spikes labeled by Kilosort2 were labeled as CS1 by P-sort (right column, Supplementary Fig. 1B). In contrast, Kilosort2 missed many CS1 complex spikes labeled by P-sort. The conditional probabilities (Fig. 10, third row) suggest a poor suppression period for both SS1 and SS2 simple spikes labeled by Kilosort2. The Kilosort2 CS1 complex spikes demonstrate an excellent suppression period, suggesting they were real. However, Kilosort2 missed a larger number of complex spikes that were labeled by P-sort and exhibited good simple spike suppression (Fig. 10, first row).

In the difficult data set, SpyKING CIRCUS produced SS labels that agreed somewhat better with P-sort than other automated software. For simple spikes, 99.3% of the SS1 labels and 93.1% of SS2 labels in P-sort were also labeled by SpyKING CIRCUS. An important difference, however, were the coincidence simple spike events, i.e., SS1 and SS2 spikes that occurred simultaneously. P-sort labeled 1640 events as coincidence of SS1 and SS2, but SpyKING CIRCUS labeled all of these as SS1 events. Another notable disagreement was among the complex spikes. While SpyKING CIRCUS found nearly the same CS1 events as P-sort (95.4% agreement), the two approaches disagreed entirely regarding the CS2 events (2.1% agreement). As the traces in the right column of Supplementary Fig. 1C illustrate, the CS2 complex spikes labeled by SpyKING are likely to be simple spikes. This conjecture appears to be confirmed by a lack of suppression in the probability traces in Fig. 10, right column, fourth row.

Because the main difference between P-sort and the automated algorithms was regarding identification of complex spikes, we thought to further verify P-sort’s performance specifically on complex spikes. We did this by comparing P-sort with a recently developed neural network (Markanday et al., 2020) that was specifically trained on complex spike waveforms. Here we found near unanimous agreement between the two approaches (Supplementary Fig. 1D). The neural network labeled 1178 CS1 and 885 CS2 events. This corresponded to 99.0% of the CS1 and 93.0% of the CS2 events labeled by P-sort. The few disagreements in the labeled events are shown on the right column of Supplementary Fig. 1D. In almost all cases, the complex spikes were preceded by a temporally adjacent simple spike, thus producing waveform distortions, making the labeling process particularly challenging.

In summary, we compared results of P-sort with automated algorithms in three data sets and found that the main difference was in the identification of complex spikes. For example, in the medium difficultly data set in which complex and simple spikes had similar waveforms, Kilosort2 missed the complex spikes. Conditional probabilities suggested that in both the easy and the very difficult data set, performance of MountainSort4 was close to P-sort, though it also could miss around 15% of the complex spikes. In the medium difficulty data set, performances of MountainSort4 and SpyKING were close to P-sort. In all three data sets, P-sort produced cleaner spike suppression periods, as well as more robust suppression of simple spikes that coincided with the labeled complex spikes. Finally, there was near unanimous agreement between P-sort and a neural network trained specifically to identify complex spikes.

### A database for testing and development of algorithms

Automated algorithms can analyze tens or hundreds of simultaneously recorded electrodes, making them essential for use with high-density probes. Unfortunately, current automated algorithms may not perform ideally in the cerebellum, as illustrated by the data here, and documented elsewhere (Hall et al., 2021). Of course, P-sort is not a solution because in its current form it relies heavily on user interaction. To facilitate development of automated algorithms for the cerebellum, we organized a large database consisting of over 300 recordings from the marmosets, macaques, and mice cerebellums. We then used P-sort to label and attribute the simple and complex spikes in each recording. Recordings in the primates were from the vermis, lobules VI and VII. Recordings from the mice were from the eye blink region of lobule V.

We found that on average, simple spike rate in the marmoset (Fig. 11A) was somewhat higher than in the macaque (63.8±1.29 Hz vs. 55.9±2.45 Hz, Mean±SEM, independent samples t-test, t(285)=2.75, p=0.006). In contrast, complex spike rate in the marmoset was somewhat lower than in the macaque (0.88±0.013 Hz vs. 0.98±0.027 Hz, t(285)= −3.6678, p= 0.0003). For the simple spikes, the conditional probability Pr(S(t) | S(0) was very similar in the two primate species (Fig. 11C), suggesting that the simple spike suppression periods are similar.

**Fig. 11.**
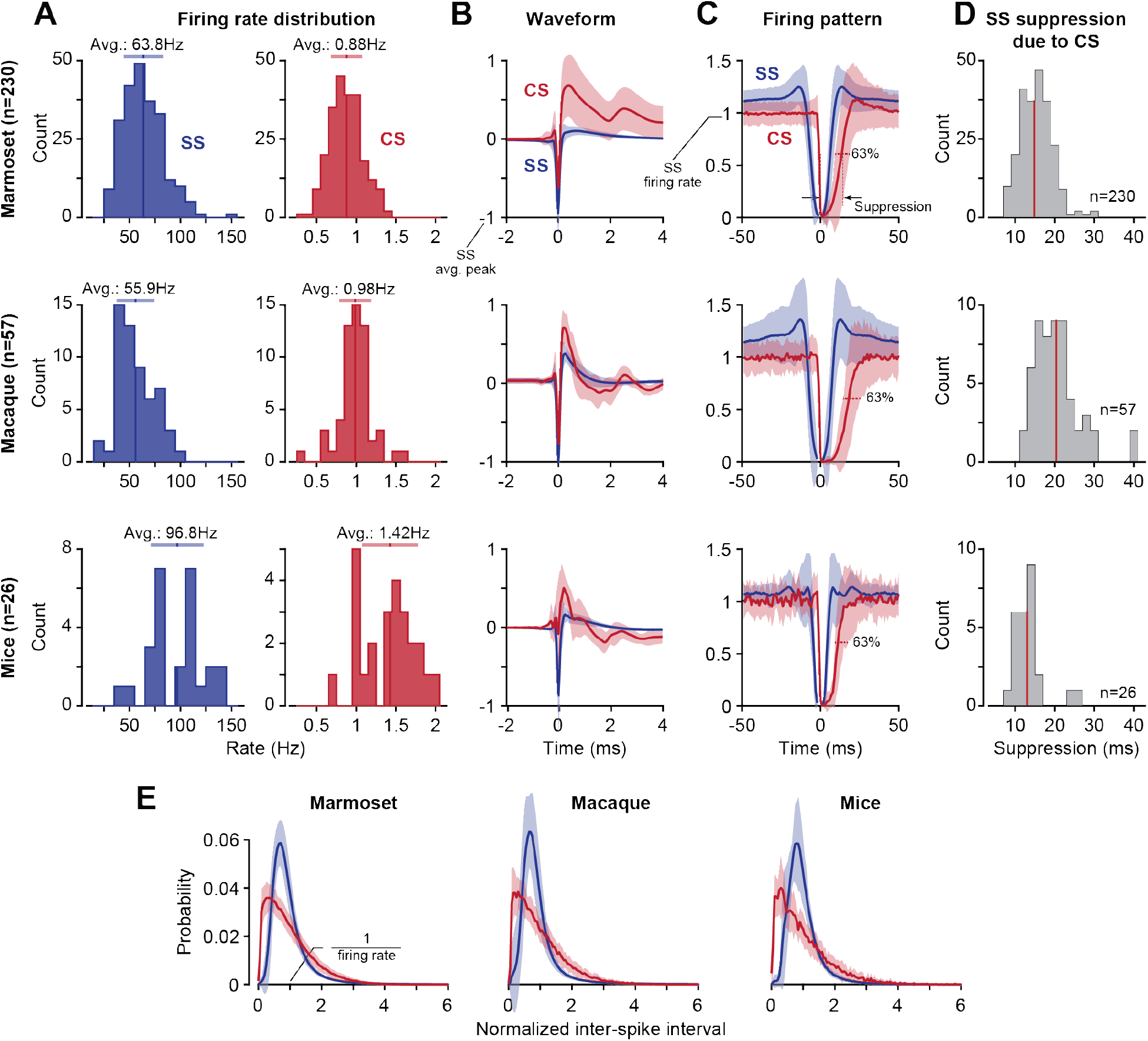
Statistical properties of simple and complex spikes in three species. **A**. Distribution of average firing rates. **B**. Waveform of simple and complex spikes. Simple and complex spikes of each P-cell were both normalized by setting to −1 the negative peak of the simple spike waveform. Error bars are standard deviation. **C**. Suppression period of simple spikes (blue, SS|SS) and the suppression coincided with complex spikes (red, SS|CS). SS|SS indicates the rate of simple spikes at time t when another simple spike occurs at time zero. SS|CS indicates the rate of simple spikes at time t when a complex spike occurs at time zero. Simple and complex spike rates for each P-cell were normalized with respect to average simple spike firing rate. Error bars are standard deviation. **D**. Suppression period of simple spikes following arrival of a complex spike. Suppression period for each P-cell was defined as the duration of time after a complex spike that was required before the simple spike rate recovered 63% of its pre-complex spike value. The red line indicates mean. **E**. Inter-spike interval distribution for simple (blue) and complex spikes (red). ISI data for each spike type in each cell was normalized so that the average ISI, defined as the inverse of the average firing rate, was equal to one. Error bars are standard deviation.

In contrast, the complex spikes in the macaque were followed by a somewhat longer period of simple spike suppression (Fig. 11C). To measure the duration of simple spike suppression that followed a complex spike, for each P-cell we computed the time *T* when the probability Pr(S(t) | C(0)) increased beyond a threshold of 63% of the baseline before onset of the complex spike (Fig. 11D). In the marmoset, P-cells had an average of 14.8±0.26 ms suppression period, significantly less than the suppression duration of 20.4±0.89 ms we quantified in the macaque (t(285)= −8.0212, p= 3e-14).

These differences are difficult to interpret because spike rates and suppression durations depend on the precise location of the P-cell. For example, P-cells located in zebrin negative regions display higher frequency simple spikes than those located in neighboring zebrin positive stripes (Zhou et al., 2014). Furthermore, the simple spike suppression following a complex spike is longer in zebrin positive zones. In both marmoset and macaque, a fraction of a millimeter difference in the recording site along the medial-lateral direction can change the zebrin band characteristics, particularly in lobule VII of the vermis (Fujita et al., 2010). Thus, the differences in rates and suppression duration between species may be due to small differences in sites of recordings.

In the mouse data the complex spike rate was somewhat higher than the marmoset, 1.42±0.07 Hz (t(254)= 12.0275, p= 1e-26), and also somewhat higher than the macaque (t(81)= 7.1277, p= 4e-10). Furthermore, the complex spikes in the mouse were followed by a shorter suppression period (12.96±0.80 ms) than in the macaque (t(81)= −5.2055, p= 1e-6) and in the marmoset (t(254)= −2.2462, p= 0.026).

Finally, in all three species we observed a consistent pattern in the distribution of inter-spike intervals (ISI). To compute the ISI distribution, we first measured the average ISI for each type of spike for each P-cell and then normalized the distribution by setting the average ISI to one. The resulting simple and complex spike normalized ISIs were different from each other, but nearly identical in the three species (Fig. 11E). These patterns may be useful priors that can aid identification of these spikes in an automated software.

In summary, we labeled simple and complex spikes in over 300 recordings in three species and quantified their statistical properties. To encourage development of automated algorithms for the cerebellum, we made the raw data as well as the P-sort labels available for 313 P-cells at https://doi.org/10.17605/osf.io/gjdm4.

## Discussion

Spike sorting in the cerebellum can be a joy, something akin to a treasure hunt: finding the correct clusters of waveforms produces a satisfying statistical pattern in which the complex spikes are followed with suppression of the simple spikes. However, identifying spikes that belong to a P-cell can be difficult both for those who prefer manual curation and those who employ automatic algorithms. While most complex spikes leave an LFP signature (Zur & Joshua, 2019), some complex spikes can lack this characteristic (Fig. 1B). Moreover, their waveforms can differ from one event to the next because of their temporal proximity to simple spikes (Markanday et al., 2020), because of spikelets (Burroughs et al., 2017; Davie et al., 2008; Ito & Simpson, 1971; Monsivais et al., 2005), or because of intrusion of spikes from neighboring neurons. Even after the spikes are identified based on their waveforms, the simple spikes may belong to one P-cell, while the complex spikes may belong to another (Fig. 1E).

To help with the identification and attribution problems, we organized a set of clustering and labeling tools in a GUI based cross-platform analysis software called P-sort. Like other sorting software, P-sort clusters spikes based on their waveform properties. However, P-sort emphasizes the statistical relationship between complex and simple spike clusters. This statistical interaction can justify merging of seemingly disparate clusters, for example when spike waveforms are distorted (Fig. 6), or justify splitting of a single cluster, for example when spikes from two different cells have similar waveforms (Fig. 7). Thus, P-sort attempts to go beyond waveform-based clustering by providing statistical information regarding how spikes interact with each other.

Our development of P-sort was aided by a diverse collection of data from marmosets, macaques, and mice cerebellums. The evaluation of the results was aided by comparison to manual curation performed by the experts in the various laboratories. In addition, we compared P-sort with automatic algorithms MountainSort4 (Chung et al., 2017), Kilosort2 (Pachitariu et al., 2016), and SpyKING CIRCUS (Yger et al., 2018), as well as a neural network trained to identify complex spikes (Markanday et al., 2020). On the easy data sets, the performances of these automated algorithms were generally good, but on medium and high difficulty data sets, they occasionally missed large groups of complex spikes, or mis-labeled them as simple spikes. A comparison of P-sort selected complex spikes with a neural network trained to identify complex spikes produced near unanimous agreement.

P-sort follows the traditional approach in which features of the spike waveforms are identified and then clustered. It relies on a recently developed algorithm called UMAP (McInnes et al., 2018), a nonlinear dimensionality reduction technique that, in our experience, is particularly powerful for clustering waveforms. UMAP has certain advantages over other non-linear dimensionality reduction algorithms such as t-SNE (t-distributed stochastic neighborhood embedding) (van der Maaten & Hinton, 2008). Unlike t-SNE, UMAP returns an invertible transform onto which new data can be projected without having to re-compute the map. This has the unique advantage of allowing for cross validation, which can be employed by semi-supervised learning methods in which the expert labels a subset of the data and leaves it to UMAP to make predictions on the unlabeled data set. Indeed, UMAP was recently adopted by Markanday et al. (2020) to cluster complex spikes from the cerebellum, and by Lee et al. (2021) to cluster spikes from the cerebral cortex.

However, UMAP has a number of disadvantages. In UMAP space, cluster sizes and distances between them do not contain information regarding the waveform structures; this issue can be resolved by viewing clusters in the PCA space, which is also provided in P-sort. Moreover, UMAP relies on a stochastic optimization process that produces non-deterministic outcomes of different runs. To help with this problem, P-sort supports a GPU implementation of UMAP that provides results for a typical 5 minute recording session in around 1-2 seconds. This rapid response makes it possible for the user to evaluate the same data set multiple times, perhaps with different waveform window sizes. In addition, in most cases UMAP can reproduce the same number of clusters with similar topological relationships. Thus, although UMAP is not deterministic, it can extract clusters that are reproducible. Regardless, there remains a possibility of overfitting in noisy recording scenarios. To help with this, the Cluster Module depicts waveforms of each cluster, as well as their statistical relationship to other spike clusters.

Once the clusters are identified, P-sort provides automated tools to find cluster boundaries. One way to improve this step is to use Louvain clustering, i.e., finding community of spikes that are highly inter-connected in the UMAP space. This approach was recently demonstrated by Lee et al. (2021) in sorting of cortical spikes.

Finding cluster boundaries, however, is not the only method to sorting, as illustrated by SpyKING CIRCUS (Yger et al., 2018). That work uses a template to define the centroid of each cluster, not their precise borders. In our tests, this approach produced good results on the large amplitude simple and complex spikes, but poor results on the smaller amplitude spikes, as illustrated by the conditional probabilities (Fig. 10).

The development of high-density electrodes highlights the need for automatic sorters. These sorters have addressed many issues including cluster matching between different channels, as well as multi-unit sorting. Moreover, their software features GUI-based visualization and manual curation toolboxes, thus allowing the user to post-process the results. However, the automated approaches currently lack GUIs for identification of dependent spikes, i.e., simple and complex spikes. In contrast, P-sort was designed to efficiently illustrate and interactively handle the simultaneous sorting and attribution of complex and simple spikes. The addition of features like multi-unit sorting opens the way for automatic sorters to be used as an initial starting point, followed by clustering and attribution by P-sort.

The rapidly evolving silicon probe technology makes it essential that we encourage development of automated sorters for the cerebellum. Thus, we used P-sort to label spikes in recordings made from over 300 P-cells in various species, and provide this labeled database to help software developers test and improve their algorithms. P-sort software is available at https://github.com/esedaghatnejad/psort. The labeled neurophysiological data are available at https://doi.org/10.17605/osf.io/gjdm4.

## Methods

### Subjects

Neurophysiological data were collected from two marmosets (*Callithrix Jacchus*, male and female, 350-370 g, subjects M and R, 4 years old), six rhesus macaques (*Macaca mulatta;* males; 5.0-7.4 kg, subjects B, F, K, P, S, and W), and ten mice (*Mus musculus;* male C57BL/6J mice at least 12 weeks of age, subjects N082, N083, N086, N089, T029, T052, T083, T101, T122, T124).

The marmosets were born and raised in a colony that Prof. Xiaoqin Wang has maintained at the Johns Hopkins School of Medicine since 1996. The procedures on the marmosets were evaluated and approved by Johns Hopkins University Animal Care and Use Committee in compliance with the guidelines of the United States National Institutes of Health.

The procedures on the macaques were performed in accordance with the Guide for the Care and Use of Laboratory Animals (2010) and exceeded the minimal requirements recommended by the Institute of Laboratory Animal Resources and the Association for Assessment and Accreditation of Laboratory Animal Care International. The procedures were approved by the local Animal Care and Use Committee at the University of Washington.

The procedures on the mice were approved by the Baylor College of Medicine Institutional Animal Care and Use Committee based on the guidelines of the US National Institutes of Health. The experimental mice were singly housed on a reverse light/dark cycle (8:00 lights-off to 20:00 lights-on).

### Marmoset data acquisition

Following recovery from head-post implantation surgery, the monkeys were trained to make saccades to visual targets and rewarded with a mixture of applesauce and lab diet (Sedaghat-Nejad et al., 2019). Using MRI and CT imaging data for each animal, we designed an alignment system that defined trajectories from the burr hole to various locations in the cerebellar vermis, including points in lobule VI and VII. We used a piezoelectric, high precision microdrive (0.5 micron resolution) with an integrated absolute encoder (M3-LA-3.4-15 Linear smart stage, New Scale Technologies) to advance the electrode.

We recorded from the cerebellum using three types of electrodes: quartz insulated 4 fiber (tetrode) or 7 fiber (heptode) metal core (platinum/tungsten 95/05) electrodes (Thomas Recording), and 64 contact high density silicon probes (Cambridge Neurotech). We connected each electrode to a 32 or 64 channel head stage amplifier and digitizer (Intan Technologies, USA), and then connected the head stage to a communication system (RHD2000, Intan Technologies, USA). Data were sampled at 30 kHz and band-pass filtered (2.5 - 7.6k Hz). We computed a common average reference signal (median of all simultaneously recorded channels, computed at 30 kHz) and subtracted this signal from each channel. We used OpenEphys (Siegle et al., 2017), an open-source extracellular electrophysiology data acquisition software, for interfacing with the RHD2000 system and recording of signals.

### Macaque data acquisition

The data were collected during previous studies (Kojima et al., 2010a, 2010b; Soetedjo et al., 2008). Following recovery from surgery, the monkeys were trained to make saccades to visual targets and rewarded with applesauce. A recording chamber was implanted on the midline of the cerebellum (14.5mm posterior of the interaural axis and directed straight down), providing access to the oculomotor vermis (lobule VI and VII). Single-unit activity was recorded with homemade tungsten electrodes with an iron-particle coating (100 kΩ impedance at 1 kHz). Neurophysiology data was sampled at 50 kHz by a Power 1401 digitizer (Cambridge Electronic Design, Cambridge, UK) and subsequently band-pass filtered (30 - 10k Hz). Data were displayed in real-time on a computer monitor running Spike2 and saved for offline analysis (Soetedjo & Fuchs, 2006).

### Mice data acquisition

Single-unit extracellular recording was performed as previously described (Heiney et al., 2018). In brief, a 2-3 mm diameter craniotomy was opened over the right side of the cerebellum (6.5 mm posterior and 2.0 mm lateral from bregma) to access lobule V and the eyeblink microzone, and the dura was protected by a layer of Kwik-Sil (WPI). A custom 3D printed recording chamber and interlocking lid (NeuroNexus) was secured over the craniotomy with dental acrylic to provide additional protection. After 5 days of recovery, the mouse was fixed in place on a treadmill via a previously implanted headplate, and Purkinje cell simple spikes and complex spikes were isolated using a tetrode (Thomas Recording, AN000968) acutely driven into the cerebellar cortex with microdrives mounted on a stereotactic frame (Narishige MMO-220A and SMM-100). The voltage signal was acquired at a 24,414-Hz sampling rate, and band-pass filtered between 100 - 10k Hz (AP channel) and between 2 - 300 Hz (LFP channel) using an integrated Tucker-Davis Technologies and MATLAB system (TDT RZ5, medusa, RPVdsEx) running custom code (github.com/blinklab/neuroblinks). The data include Purkinje cells from a previously collected dataset (Achilly et al., 2020; Ohmae & Medina, 2015).

### P-sort main window

To allow P-sort to run on Windows, MacOS, and Linux, the code was written using Python-based libraries (Behnel et al., 2011; Harris et al., 2020; Pedregosa et al., 2011; Virtanen et al., 2020). The GUI was written using PyQt5 (The Qt Company and Riverbank Computing Ltd.) and PyQtGraph to provide a fast and intuitive interaction for the user. To facilitate further development of P-sort by the user community, we used object-oriented coding. P-sort’s source code is available for download at: https://github.com/esedaghatnejad/psort.

A process starts by loading the data and dividing it into one or more periods of time (called slots). The slot framework helps the user to account for potential drift and fluctuation in spike quality and shape over time. After sorting one slot, the parameters and waveform templates will be copied to the next slot to facilitate the sorting, but the user can further change the parameters independently in each slot.

The sorting process starts by filtering the signal into two channels, Action Potential (AP) and Local Field Potential (LFP). The default is a 4^th^ order Butterworth filter with the 50-5000 and 10-200 Hz range for AP and LFP channels, respectively. However, these parameters can be modified using the GUI to better fit the specific conditions of the data. The default assumption is that simple spikes generate negative peaks in AP channel and complex spikes generate positive peaks in the LFP channel. However, this assumption can be changed via the GUI. Once the respective peaks are detected, the next question is what should be the threshold to reject or accept a peak as being a potential spike. P-sort computes the histogram of the peaks and fits a Gaussian Mixture Model (GMM) with two basis functions to the histogram for each channel. The lower bound Gaussian is considered the noise and the upper bound Gaussian is the signal of interest. Based on this assumption, the intersection of the two fitted Gaussians is used as the default threshold to prune the detected peaks. However, the user is provided with a GUI to manually change the SS and CS thresholds, either by using the interactive plots, or directly by assigning their values.

The next question is how to relate a peak in the LFP channel, which may potentially be a complex spike, to its waveform in the AP channel. In a typical recording, a CS waveform consists of an initial sharp negative peak and a broad positive bump. The peak in LFP happens due to the broad positive bump but its timing is variable. Thus, using the LFP peak to align the CS waveform is unreliable. P-sort provides three different methods to align complex spikes: SS index, SS template, and CS template. Initially, it uses the detected sharp negative peak (SS index) to align CS waveforms. This provides a reliable set of waveforms to calculate the CS template. However, due to variability in CS waveforms, not all CSs express the initial sharp negative peak. To address this problem, after forming a CS template, that template will be used to align CS waveforms. For the alignment, we move the 3.5 ms template signal along 1 ms past and 4ms before the LFP peak on the AP signal and select the point of time which results in maximum correlation between the two signals. For the recordings in which the LFP peak is later than 4 ms after the sharp negative peak, this default value should be adjusted using the *Preferences* interface. Alignment of the simple spikes relies on the timing of the peak value of the waveform.

P-sort ensures that a candidate spike is labeled as either a simple spike, or a complex spike, but not both. Moreover, due to biological refractory period in a spiking neuron, two arbitrary spikes cannot happen closer than 0.5 ms with respect to each other. Based on these constraints, P-sort addresses potential conflicts between CS-CS, CS-SS, and SS-SS candid events. The default values for each scenario can be changed using the *Preferences* interface.

After resolving the potential conflicts, P-sort provides sets of potential simple and complex spikes. For each set of spikes, P-sort represents spike waveforms, instant-firing-rate distribution, peak distribution, conditional probabilities, and feature scatter plots. Numerous features can be used for clustering of these data, including UMAP, principal components, timing of the spikes, relative time with respect to next or previous spike, similarity to templates, peak value, and instantaneous firing rate. Using the interactive plots, the user can select subset of the spikes based on the waveform plot or feature scatter plot and further prune the data or even change their label from simple to complex spikes and vice versa. As these clusters are manipulated, P-sort provides real-time feedback on their statistical features, thus allowing the user to determine whether the simple and complex spikes are likely to have been generated by a single P-cell, and whether the latest manipulations of the clusters improved the probabilities.

Overall, P-sort’s main window aims to provide a balance between the ability to visualize each spike waveform, and the ability to cluster the spikes and visualize their interactions. From this main window P-sort branches into two additional windows: the *Cluster Module*, and the *Dissect Module*.

### Cluster Module

A unique challenge in cerebellar neurophysiology is finding the simple and complex spike clusters that belong to a single P-cell. It is possible that on certain recordings, one or more neurons will contribute to the signals that are recorded by a single contact. For example, it is possible that the large amplitude simple spikes are not produced by the P-cell that has produced the complex spikes in the recording (Fig. 1D). Rather, the smaller amplitude simple spikes should be attributed to the complex spikes.

The Cluster Module gives the user the ability to assign labels to each waveform cluster (or part of a cluster), and immediately assess the statistical interactions between the labels. To use this module, the user will select the spikes of interest either from the feature scatter plot or the waveform plot and assign a label to the selection. Cluster Module provides four interactive subpanels for representing (1) color-coded feature scatter plot of spikes, (2) spike waveforms of each label, (3) peak histogram of each label including the firing rates information, and (4) cross and auto correlograms of chosen labels.

For example, let us assume that the user has assigned SS-1, SS-2, and CS-1 cluster labels. Then, the user can address the attribution problem of potential simple spikes with the candidate complex spike by checking the correlogram plots.

Cluster Module toolbox includes manual and automatic labelling tools to label datasets based on their features or waveforms. The automatic algorithms implemented for clustering include: (1) Gaussian Mixture Models (GMM), which requires the user to specify the number of clusters and their initial centroids, (2) Iso-split algorithm (Chung et al., 2017; Magland & Barnett, 2016), which is automatic and determines the number of clusters based on bimodality tests, and (3) HDBSCAN algorithm (Campello et al., 2013), which is also automatic and requires no user inputs. We implemented a post-processing layer for HDBSCAN’s outputs and restricted the number of clusters to less than 10. We did this by setting extra clusters with least number of members as noise (assigned as label −1). In addition to automatic clustering algorithms, an outlier detection method was implemented based on Local Outlier Factor density (Pedregosa et al., 2011) which receives the quantile threshold as input. All these algorithms use the selected elements of the feature scatter plot by default; however, the user can perform multi-dimensional clustering by selecting the multiple features from the feature list in the GUI.

### Dissect Module

P-sort dissect module is designed to provide more tools for reviewing individual spikes. In some scenarios, looking at the individual spikes and their surrounding events provides a better insight than the average features. Dissect module provide a tool set to move between spike events and look at each one over time. This module also provides the tool to manually overwrite a spike or change its alignment.

### Comparison with expert curation

50 sessions (34 in the macaque and 16 in mouse) were sorted using Spike2 and P-sort by different experts. For Spike2 data, we ensured that a given spike was not labeled as both a CS and a SS and removed the overlapping labels. Next, we compared P-sort data with Spike2 data by finding shared complex/simple spikes that happened in a 0.5/5.0 ms window of each other. We used the window of time to account for the variability in alignments of Spike2 data due to lack of template matching. If a given spike was not shared between the two datasets, it was labeled as an exclusive spike. In order to compare the number of exclusive spikes over datasets, we normalized the number of the exclusive spikes by the total number of the spikes in each dataset. We considered the Spike2 and P-sort results as separate datasets and other than finding the shared spikes, we did not crossed the results.

### Comparison with automatic sorters

We quantified performance of various automatic sorting algorithms on three different cerebellar data, ranging from easy to medium to hard. The hard data set was named the “*P-sort challenge*”. Each dataset included around 15 minutes of recording from the marmoset oculomotor vermis, and contained some of the challenges that are present in cerebellar neurophysiology, including diverse patterns of spikes from neighboring cells. Thus, sorting of the data required isolating different spike types, as well as addressing the CS-SS attribution problems. The same 0.5/5.0 ms window was used to detect shared and exclusive spikes between automatic sorters and P-sort data.

For automatic sorters, P. Yger sorted the data using SpyKING CIRCUS (Yger et al., 2018), and A. Markanday sorted the data using a neural network (Markanday et al., 2020). In addition, we used Mountainsort4 (Chung et al., 2017) and KiloSort2 (Pachitariu et al., 2016). We post-processed the outputs by merging and associating simple spikes, complex spikes, and multi-unit activity (MUA) based on rates and cross correlograms to arrive at the best match. For Mountainsort4, we used the default parameters and then the output units were manually merged and labelled as SS1, SS2, CS1, CS2, and MUA. Similarly, for Kilosort2, we used the default parameters and manually merged and labeled the outputs using the Phy2-interface correlograms and rates.

## Acknowledgements

The work was supported by grants from the National Science Foundation (CNS-1714623), the NIH (R01-EB028156, R01-NS078311, R01-EY028902, R01-EY023277, P51-OD010425, R34-NS118445, R01-MH093727, RF1-MH114269, R01-NS112917), and the Office of Naval Research (N00014-15-1-2312).

---

**Supplementary Fig. 1.**
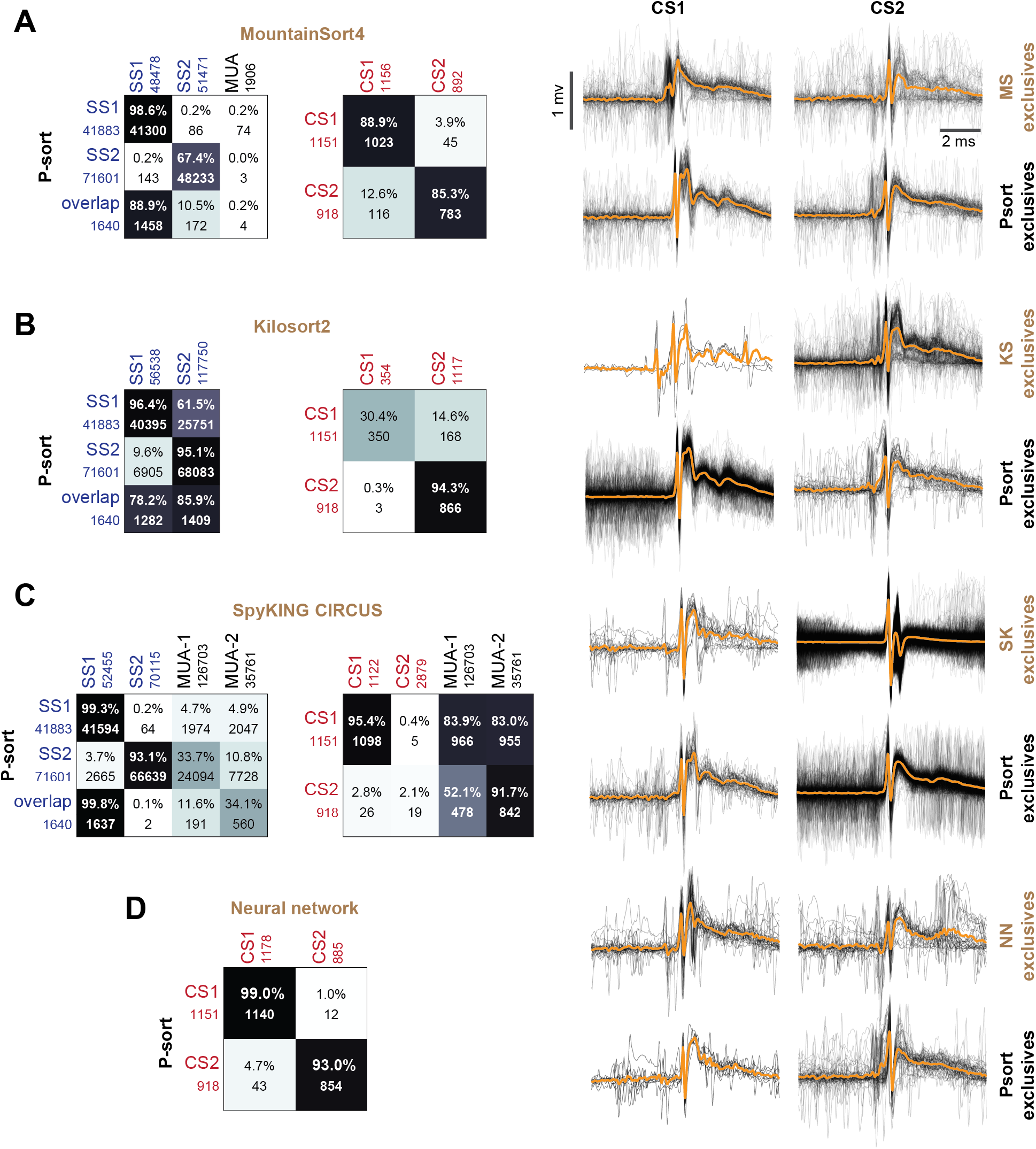
Comparison of P-sort with automated spike sorting algorithms on a difficult data set (P-sort challenge data set). The data are from a marmoset recording that contained multiple clusters of simple and complex spikes (the same data were presented in Fig. 6). **A**. Comparison to MountainSort4. The right column shows complex spikes CS1 and CS2 that were labeled exclusively by each algorithm. **B**. Kilosort2 missed roughly 70% of the CS1 complex spikes labeled by P-sort. **C.** SpyKING CIRCUS disagreed entirely with P-sort regarding complex spike CS2 cluster. **D**. Comparison of P-sort complex spike identification with a neural network trained to identify complex spikes (Markanday et al., 2020).

